# Co-expression of Mitochondrial Genes and ACE2 in Cornea Involved in COVID-19 Infection

**DOI:** 10.1101/2020.07.23.216770

**Authors:** Jian Yuan, Dandan Fan, Zhengbo Xue, Jia Qu, Jianzhong Su

**Author notes:** These authors contributed equally to the paper as first authors (Jian Yuan, Dandan Fan).

## Abstract

The Coronavirus disease 2019 (COVID-19) pandemic severely challenges public health and necessitates the need for increasing our understanding of COVID-19 pathogenesis, especially host factors facilitating virus infection and propagation. Here, the co-expression network was constructed by mapping the well-known ACE2, TMPRSS2 and host susceptibility genes implicated in COVID-19 GWAS onto a cornea, retinal pigment epithelium and lung. We found a significant co-expression module of these genes in the cornea, revealing that cornea is potential extra-respiratory entry portal of SARS-CoV-2. Strikingly, both co-expression and interaction networks show a significant enrichment in mitochondrial function, which are the hub of cellular oxidative homeostasis, inflammation and innate immune response. We identified a corneal mitochondrial susceptibility module (CMSM) of 14 mitochondrial genes by integrating ACE2 co-expression cluster and SARS-CoV-2 interactome. Gene ECSIT, as a cytosolic adaptor protein involved in inflammatory responses, exhibits the strongest correlation with ACE2 in CMSM, which has shown to be an important risk factor for SARS-CoV-2 infection and prognosis. Our co-expression and protein interaction network analysis uncover that the mitochondrial function related genes in cornea contribute to the dissection of COVID-19 susceptibility and potential therapeutic interventions.

## Introduction

A novel coronavirus, severe acute respiratory syndrome coronavirus 2 (SARS-CoV-2) associated with severe human infected disease (COVID-19) outbreak starting from December 2019, in China, and the disease is quickly spreading worldwide (1). Despite being primarily a respiratory virus, COVID-19 can also present with non-respiratory signs, including ocular symptoms as conjunctival hyperemia, chemosis, epiphora, increased secretions, ocular pain, photophobia and dry eye (2). The presence of virus in tear, conjunctival swab specimens and animal models of infectious increasing clinical and scientific evidence that eye may serve as a potential site of virus replication (3, 4). Moreover, immunohistochemical studies and single-cell RNA-sequencing datasets revealed both extra- and intra-ocular localisation of SARS-CoV-2 entry factors, ACE receptor and TMPRSS2 protease in human eyes (5, 6). Together, these results suggest that ocular surface cells are susceptible to infection by SARS-CoV-2. More a recent GWAS study identified 3p21.31 as a most significant genetic locus being associated with COVID-19 induced respiratory failure (7). This locus covers a cluster of six genes consisting of SLC6A20, LZTFL1, CCR9, FYCO1, CXCR6, and XCR1, with the identified risk allele being associated with increased SLC6A20 and LZTFL1 expression. Of note, SLC6A20, LZTFL and FYCO1 is known to associate with eye development, electroretinography abnormal and anterior eye segment morphology. However, whether these key factors for cellular susceptibility to viral infection have correlation in ocular surface cell remains unclear. Herein, we constructed co-expression and interactome networks and mapped genes implicated by COVID-19 GWAS onto corneal co-expression network to infer the function of susceptibility gene.

## Results

To understand the expression patterns of ACE2, TMPRSS2 and susceptibility genes in cornea, we firstly compared the expression level of ACE2 in cornea, retina, RPE and lung tissues based on bulk RNA sequencing. As expected from prior literature (8), the cornea showed a higher ACE2 expression than lung both in terms of their TPM values (Fig 1A). ACE2 exhibits the highest co-expression correlation with TMPRSS2, SLC6A20C and LZTFL1 in the cornea compared to lung and RPE (Fig. 1B). To gain more insight into the biological network of genes associated with SARS-CoV-2 entry factors and susceptibility gene, we performed k-means clustering algorithm to identify genes associated with ACE2 on cornea datasets (Fig. 1C). We identified 26 co-expression modules ranging in size from 111 to 1798 genes. One cluster contained both ACE2, LZTFL1 and FYCO1 (cluster 7; 1434 genes). The strongest correlation with the eigengene (the principal component) of this ACE2 cluster was found for hub genes STK16 (r=0.95, p=1.07×10^−9^), a member of NAK family that activate the AP□2 scaffolding protein vital to viral entry and propagation. TMPRSS2 belonged to a separate cluster 5 (603 genes) with hub gene SLC25A1 (r=0.91, p=4.45×10^−8^), which involved in TNF-α and IFN-γ triggered inflammation. Together these data suggest that the cornea may provide a susceptibility and entry portal for the SARS-CoV-2 entry.

**Figure 1.**
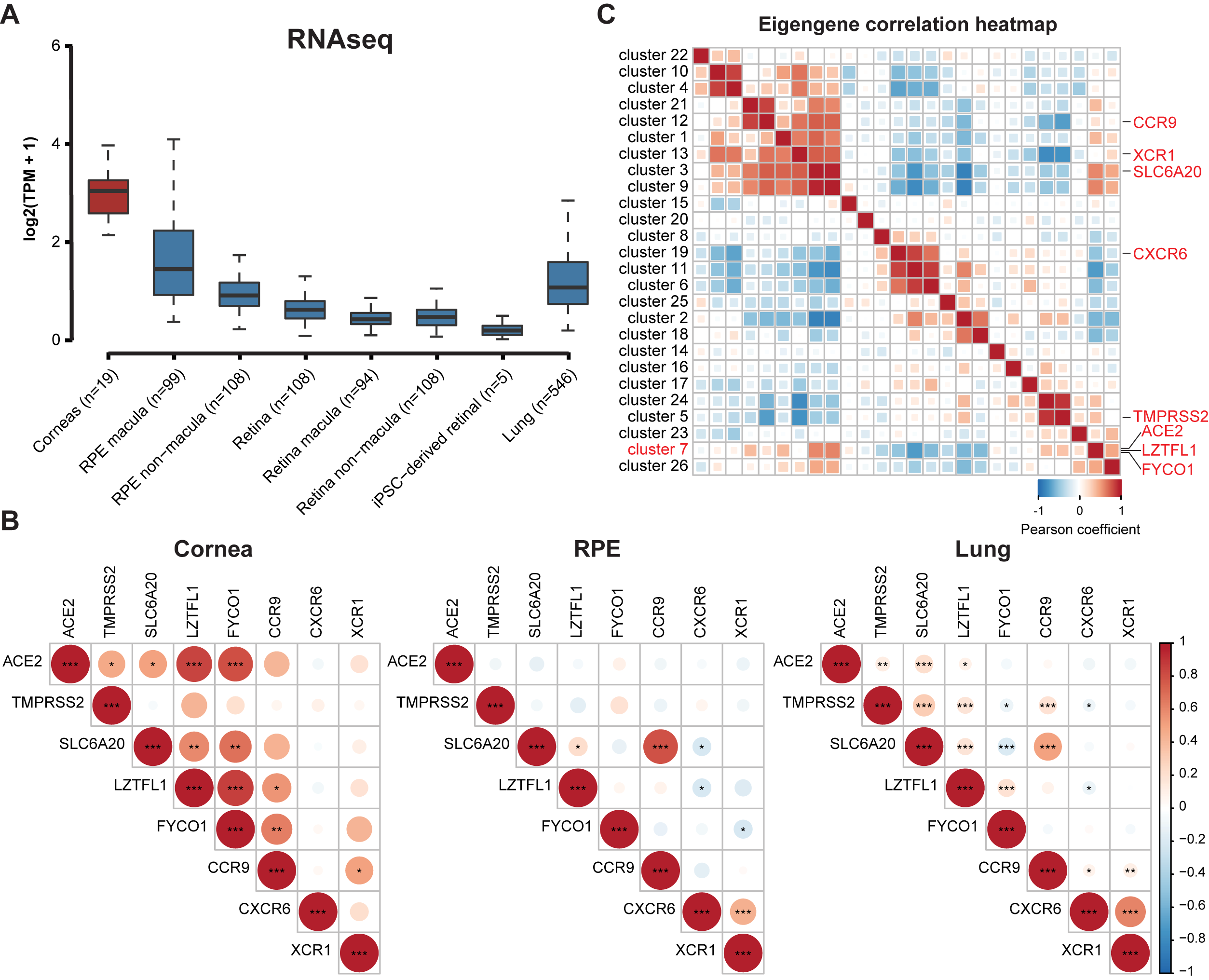
Expression of ACE2 across different eye tissues and its co-expression pattern in the cornea. A. Expression of ACE2 in cornea, retina, RPE and lung. B. Pearson’s correlation analysis were estimated between ACE2 and TMPRSS2 and six susceptibility genes. The size of the circle scales with the correlation magnitude. The darker the color, the larger the magnitude of correlation coefficient. Star sign (*) indicates statistical significance. C. Eigengene correlation heatmap representing the strength and significance of correlations between cluster eigengenes. Pearson’s correlation coefficient is used as the correlation descriptor (red and blue for positive and negative correlations, respectively).

To gain insights into the molecular functionality related to ACE2 cluster, we integrated the ACE2 cluster and SARS-CoV-2 interactome and observed their shared a number of similarities. For instance, the ACE2 cluster and SARS-CoV-2 interactome eigengenes were highly correlated (r=0.56, p=0.014). Genes within ACE2 cluster were mainly related to mitochondrion functions such as mitochondrial inner membrane and mitochondrial electron transport (p=7.47×10^−29^) (Fig. 2A). We further evaluated shared GO enrichments to determine whether the similarity in the behavior meant that both clusters contained functionally related genes. ACE2 cluster and SARS-CoV-2 interactome were nominally enriched (p value <0.001) for 41 and 35 GO terms, respectively. Of these, 17 terms were enriched in both modules. Furthermore, we observed a positive correlation in fold enrichments for shared terms (Fig. 2B). Most of the ontologies shared between the modules described cellular components, biological processes, and molecular functions pertinent to mitochondrion (Fig. 2B). In both the ACE2 cluster and SARS-CoV-2 interactome, the observed numbers of mitochondrial genes were significantly higher than the number what would be expected by chance (p=2.1×10^−14^ and p=3.2×10^−6^, respectively; Fig. 2C). In addition, we found the observed number of ACE2 cluster that was significantly higher (p = 0.005) associated with interaction protein of SARS-CoV-2 compared to randomness (Fig. 2C). Together, these results indicated that ACE2 cluster possess the key factors required for cellular susceptibility to SARS-CoV-2 infection in cornea. Therefore, the core 14 genes were co-occurred in all three datasets as the corneal mitochondrial susceptibility module (CMSM) of SARS-CoV-2 infection in cornea (Fig. 2D). Of the 14 CMSM genes, five genes have been shown to directly implicate in the function of electron transport. They include genes such as NDUFB9 a member of mitochondrial respiratory-chain complex 1, and NDUFAF1, NDUFAF2, ECSIT which are involved in mitochondrial respiratory-chain complex 1 assembly. Next, we studied the expression alteration of these genes in cornea from 23 keratoconus patients and 19 health controls. We found that ACE2 expression was significantly increased in keratoconus compared to control cornea (log_2_FC=2.8, p=4.4×10^−7^; Fig. 2E). Other than the up-regulation of ACE2 in keratoconus, there were 10 of 21 genes related to SARS-CoV-2 infection was significantly upregulated in keratoconus patients (Fig. 2E, t-test p-value < 0.05). Based on the elevated expression of ACE2 and other susceptibility genes in keratoconus, we speculated that patients with keratoconus are more likely to be infected by the SARS-CoV-2.

**Figure 2.**
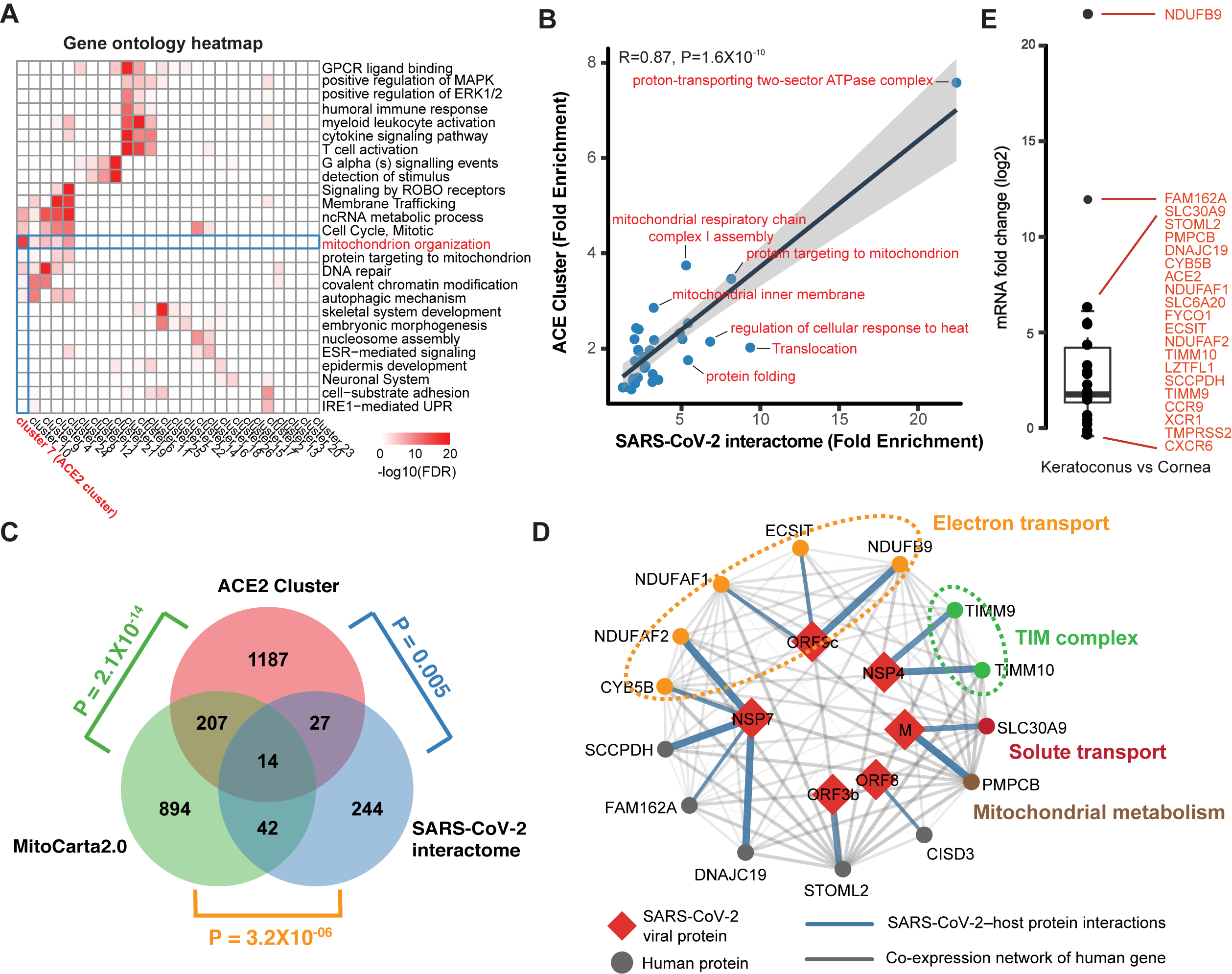
Identification of the Cornea Functional Module and Mitochondrial Susceptibility Genes. A. Clustered heat map of gene ontology (GO) terms among cluster genes. Color coding according to legend at the bottom, only gene ontology terms with FDR < 0.05 were considered. B. Gene ontology fold enrichments are correlated for GO terms shared between ACE2 cluster and SARS-CoV-2 interactome. C. Venn diagram depicting the overlap between ACE2 cluster, SARS-CoV-2 interactome and mitochondrial genes. *P*-value computed using Fisher’s exact test. D. SARS-CoV-2 protein-protein interaction between 14 mitochondrial susceptibility genes (circles) and 6 SARS-CoV-2 proteins (red diamonds). Blue edge thickness proportional to interaction MiST score; Grey edge thickness proportional to correlation of gene expression. E. mRNA abundance change of ACE2, TMPRSS2 and susceptibility genes in keratoconus.

## Discussion

To control and mitigate the impact of the COVID-19 pandemic, it is vital to gain greater understanding of the routes and modes of transmission, including the role of the ocular surface. Using a cornea-relevant co-expression network to inform virus entry factors and GWAS interpretation, we were able to identify putative susceptibility genes for highly correlated with ACE2. Interestingly, we found that ACE2 co-expression cluster and SARS-CoV-2 interactome were both enriched for processes to respiratory electron transport which are being targeted by metformin. A favourable effect of metformin in patients with COVID-19 has been hypothesised as the drug might prevent virus entry into target cells via adenosine monophosphate-activated protein kinase activation and the phosphatidylinositol-3-kinase–protein kinase B-mammalian target of rapamycin signalling pathway (9). Moreover, we predicted that the mitochondrial related CMSM gene set was vital susceptibility genes based on the co-expression and protein interaction network. Notably, ECSIT, one of the CMSC gene, is a cytosolic adaptor protein involved in inflammatory responses and plays a regulatory role as part of the TAK1-ECSIT-TRAF6 complex that is involved in the activation of NF-κB by the TLR4 signal. In the previous study, treatment with drugs that inhibited NF-κB activation led to a reduction in inflammation and significantly increased mouse survival after SARS-CoV infection (10). Additionally, ECSIT is also essential for the association of RIG-I-like receptors (RIG-I or MDA5) to VISA in innate antiviral responses. Therefore, ECSIT may be used as a new drug target to protect against the development of severe forms of COVID-19 infection. Based on our results, we believe that significant insight into COVID-19 in cornea can be gained using co-expression and interaction networks.

## Materials and Methods

We collected and analyzed the several RNA-seq studies of the normal tissue of cornea (n=19 samples) (11), retina (n=310 samples), retinal pigment epithelium (RPE) (n=207 samples) (12) and lung (n = 546 samples) from the NCBI GEO database (GSE77938 and GSE115828) and GTEx database. To specifically characterize the biological pathways involved, we performed k-means analysis, identifying several co-expression clusters. For further information about data, methods, and code, see https://github.com/DanielWYuan/COVID-19_Cornea.

## Acknowledgments

This work was supported by the Key Program of National Natural Science Foundation of China (81830027) to J. Qu; the National Natural Science Foundation of China (61871294), Zhejiang Provincial Natural Science Foundation of China (LR19C060001) to J. Su

